# Comprehensive evaluation of antibody responses to mosquitoes and mosquito-borne arboviruses using highly multiplexed serology

**DOI:** 10.1101/2025.08.15.670562

**Authors:** Melodi Anahtar, Diana Striplet, Eben Philbin, Joshua R. Lacsina, Jesus G. Valenzuela, Michael Johansson, Fabiano Oliveira, Laura Willen, Shaden Kamhawi, Auden Cote-L’Heureux, Sarah M. Fortune, Daniel Neafsey

## Abstract

Vector-borne diseases (VBDs) are a leading cause of morbidity and mortality worldwide, and their future impact may increase due to climate change. Antigens driving host immune responses to pathogens and vectors can serve as vaccine candidates and biomarkers of previous exposure, but the immunogenicity of the relevant proteomes remains undercharacterized. To comprehensively profile antibody responses to VBDs and to the vectors themselves, we developed a highly multiplexed phage display library (VectorScan) containing over 250,000 peptides derived from diverse arthropod vectors and prevalent vector-borne pathogens. We used phage immunoprecipitation sequencing (PhIP-Seq) to screen VectorScan against blood samples from non-human primates and humans with experimental and natural exposures to arboviruses and mosquitoes (*Anopheles gambiae* and *Aedes aegypti*). We analyzed quantitative measurements of peptide seroreactivity to identify epitopes driving viral serotype-level exposure signatures, as well as novel mosquito sialome antigens. Mosquito-directed antibody responses were associated with natural viral exposure, but highly heterogeneous at an individual level. However, recurrent responses to insect-derived cuticular proteins, mucin-like proteins, and fibrinogen-like proteins will inform development of future serosurveillance tools for vector exposure and vector-based vaccines.

## Introduction

Each year, vector-borne diseases (VBDs) cause over 700,000 deaths and hundreds of millions of infections around the world.^1^ Arthropod vectors such as mosquitoes and ticks accomplish pathogen transmission through blood feeding behavior. During feeding, an arthropod not only ingests blood, but also injects saliva and any pathogens, including those present in the salivary glands. The proteins in saliva, referred to collectively as the sialome, can enhance pathogen transmission and increase mortality in animal models of infection compared to needle-based inoculation, through mechanisms such as host immune activation and immune cell recruitment to the bite site.^2^ This has motivated the exploration of the use of immunogenic salivary proteins as anti-vector vaccines.^3^ While the immunogenicity of vector sialomes has not been comprehensively assessed, vaccination with candidate sialome antigens has been shown to prevent infection in animal models of major VBDs including Lyme, Zika, and malaria.^4–6^ The immunogenicity of salivary components has also motivated their use as population-level biomarkers of vector exposure via serosurveillance.^7,8^ However, immune responses to individual sialome antigens assayed via ELISAs are too variable to reliably infer individual-level exposure.^9,10^ Ultimately, the ability to detect serological biomarkers of exposure to vector-borne pathogens and their vectors simultaneously could be used to connect vector exposure with infection risk, inform public health measures, identify shifts in vector ranges and disease incidence with climate change, train ecological models, and determine new vaccine targets for VBDs. We therefore developed an exploratory phage display library called VectorScan to broadly profile the immunogenicity of diverse vector sialomes and simultaneously measure exposure to VBDs.

Phage display libraries enable broad surveys of immunogenicity by encoding small peptides into bacteriophages that express the protein on their surface, associating each encoded DNA sequence with the properties of its expressed peptide. These libraries can be screened using phage immunoprecipitation sequencing (PhIP-Seq) to simultaneously interrogate antibody responses to hundreds of thousands of peptides 25-60 amino acids in length.^11,12^ These peptides can be designed to tile through the potentially antigenic protein sequences of an organism or infectious agent of interest, creating a library of potential linear epitopes to which host-derived antibodies generated against these antigens can bind. Several phage display libraries have been previously developed to query antibody responses to vector-borne pathogens. These include “ArboScan”,^13^ which is focused on arboviruses, the “Falciparome”,^14^ which is focused on the *Plasmodium falciparum* malaria parasite,^15^ and smaller yeast display libraries designed to query the humoral response to tick and mosquito antigens.^16,17^

To broadly measure individual exposure to diverse vector-borne viral, bacterial, and parasitic pathogens, as well as exposure to the sialomes of diverse arthropod vectors, we developed a phage display library called VectorScan, the first version of which incorporates ∼250K peptides from 15 pathogens and 24 vectors. In this work, we focus on mosquito antigens and mosquito-transmitted arboviruses that were included in the library. We used VectorScan to detect mosquito- and dengue-derived antibodies in animal and human blood samples with known and controlled exposure histories. We also profiled anti-mosquito and anti-arbovirus antibody repertoires among field samples collected from individuals with and without IgG seropositivity to arboviruses. We anticipate that the ability to quantitatively and simultaneously query both anti-vector and anti-pathogen responses using the same serological tool will help increase the throughput with which researchers uncover immunogenic targets for vector-borne diseases and make connections between vector exposure and infection risk.

## Methods

### Sample sourcing

We sourced sera via the BEI Resources Repository from three non-human primates (NHPs) that were each exposed to a different dengue virus strain from distinct serotypes (NHP1: infected with dengue virus 1 Nauru/West Pac/1974 [DENV1]; NHP2: infected with dengue virus 2 Thailand/NGS-C/1944 [“DENV2”]; NHP4: infected with dengue virus 4 Dominica/814669/1981 [“DENV4”]). “Pre-exposed” samples were collected prior to infection, “Early-immune” sera were collected 30 days post-exposure, and “Late-immune” sera were collected monthly from 120 to 390 days post-exposure. A pre-exposed sample was unavailable for DENV3, so it was not included in this study. Human plasma samples from controlled mosquito exposures conducted in NCT03641339 were de-identified, transferred from the National Institutes of Health (NIH) to Harvard University under a material transfer agreement, and approved for use under human subjects protocols IRB19-1317 and IRB22-1649.^18^ These samples were from healthy individuals exposed to either *Anopheles gambiae* or *Aedes aegypti* mosquitoes four times over the course of six weeks (the Cohort B study arm). Samples were collected before feedings (Day 0) and again after all four exposures (Day 44). The Centers for Disease Control (CDC) Dengue Branch in Puerto Rico provided a set of de-identified human plasma samples for VectorScan evaluation. These samples were collected prior to 2019 from individuals who were not sick at the time of sample collection and were tested by ELISA for anti-DENV IgG and anti-CHIKV IgG, and were screened with VectorScan under the same human subjects protocols.

### Design of the oligo library

Because the experiments in this work are focused on mosquitoes, we highlight the design of the mosquito-relevant portions of the library. However, detailed information about other sections of the library (i.e., ticks and *Plasmodium*) can be found in the **Supplementary Methods** and accompanying code. Briefly, we selected 21 mosquito species (species information in **Table S1**) from across the *Aedes, Anopheles*, and *Culex* genera to include in this pilot. We downloaded 378,082 protein sequences from UniProt representing the complete proteomes of these mosquito species. We retained a subset of these proteins for library peptide design based on homology to five well-annotated salivary transcriptomes from five key vector species: *Aedes aegypti*,^19^ *Anopheles gambiae*,^20^ *Culex quinquefasciatus*,^21^ *Ixodes scapularis*,^22^ and *Xenopsylla cheopis*.^23^ Further filtering to eliminate redundant proteins with excessively homologous sequence and absence of predicted secretion using SignalP narrowed the list to 156,876 mosquito-derived peptides (**Table S1**).^24^ To design the arbovirus section of the library, we downloaded all available UniProt sequences from 12 arboviruses yielding 51,136 protein sequences (species information in **Table S1**), which was reduced to a list of 2,016 proteins after clustering based on a threshold of 98% identity. We tiled through these sequences in their entirety, ultimately resulting in 18,538 arbovirus-derived peptides (**Table S1**). We also included peptides containing random sequences to act as negative controls, and another set of peptides from common viruses (such as Rhinovirus B) to serve as positive controls. The *pepsyn* package was then used to perform reverse translation of each peptide into a DNA sequence, add sequence adapters containing EcoRI and XhoI digest sites (5’-TGAATTCGGAGCGGT-3’, and 5’-CACTGCACTCGAGA-3’) to each end of the resulting oligo, and re-code any potential cleavage sites within the DNA sequences.^25^ The final oligo library (253,789 peptides total) was then synthesized by Twist Bioscience.

### Phage Immunoprecipitation Sequencing (PhIP-Seq)

The oligo phage library was cloned into T7 bacteriophage, packaged, amplified, and used for PhIP-Seq using a slightly modified version of previous protocols.^26,27^ Details of the PhIP-Seq protocol are in the **Supplementary Methods**, but briefly, we incubated the library with serum or plasma, allowing the antibodies in the sample to bind to the phage expressing immunogenic epitopes. The antibody-phage complexes were pulled down using magnetic beads that bind to IgG, amplified to increase the detectable signal, sequenced, and demultiplexed to determine the counts of each recognized peptide per sample. After sequencing, the resulting peptide counts were first normalized by sequencing depth within each replicate (based on read counts per 100,000 reads, RPK). We then calculated the fold change between the RPK of each sample (treating replicates separately) over the median RPK across the no-serum controls on a per-peptide basis. The resulting normalized fold change values are reflective of both the relative abundance of each peptide within a sample and the relative level of signal above what would be expected by nonspecific binding.

In this work we use specific criteria and terminology to characterize peptide seroreactivity. A peptide is considered “seroreactive” if both replicates from a sample are above a sample-specific noise threshold, determined by the read count distribution from the random control peptides (see **Supplementary Methods** for details). A peptide is “seronegative” if both replicates are below this threshold. Alternatively, the description of “not seroreactive” encompasses two states: either the peptide is seronegative, or it does not reach the seroreactivity threshold in both replicates.

## Results

We developed a phage library designed to simultaneously query the antibody response to the proteomes of diverse viruses, bacteria, and parasites, and the sialomes of diverse mosquito and tick vectors. We used a homology-based approach to identify a broad set of proteins from our organisms of interest to interrogate for immunogenicity (see **Methods**). The final set of 253,789 peptides (**Fig. 1A**, sequences in **Table S2**) from mosquitoes, arboviruses (spanning alphaviruses, bunyaviruses, flaviviruses), and other VBD-relevant organisms, were synthesized into an oligo library and packaged into T7 bacteriophage as previously described.^25^ VectorScan, the resulting phage library, has over 99.75% coverage of the desired sequences (sequenced to ∼250x coverage, **Fig. S1A)**, and was used to perform PhIP-Seq (**Fig. 1B**). Downstream analyses were performed on a subset of 249,243 peptides (∼98.2% of the original library), after excluding those that were not represented after expansion and those with large GS-linkers (see **Table S1** and **Supplementary Methods** for details).

**Figure 1:**
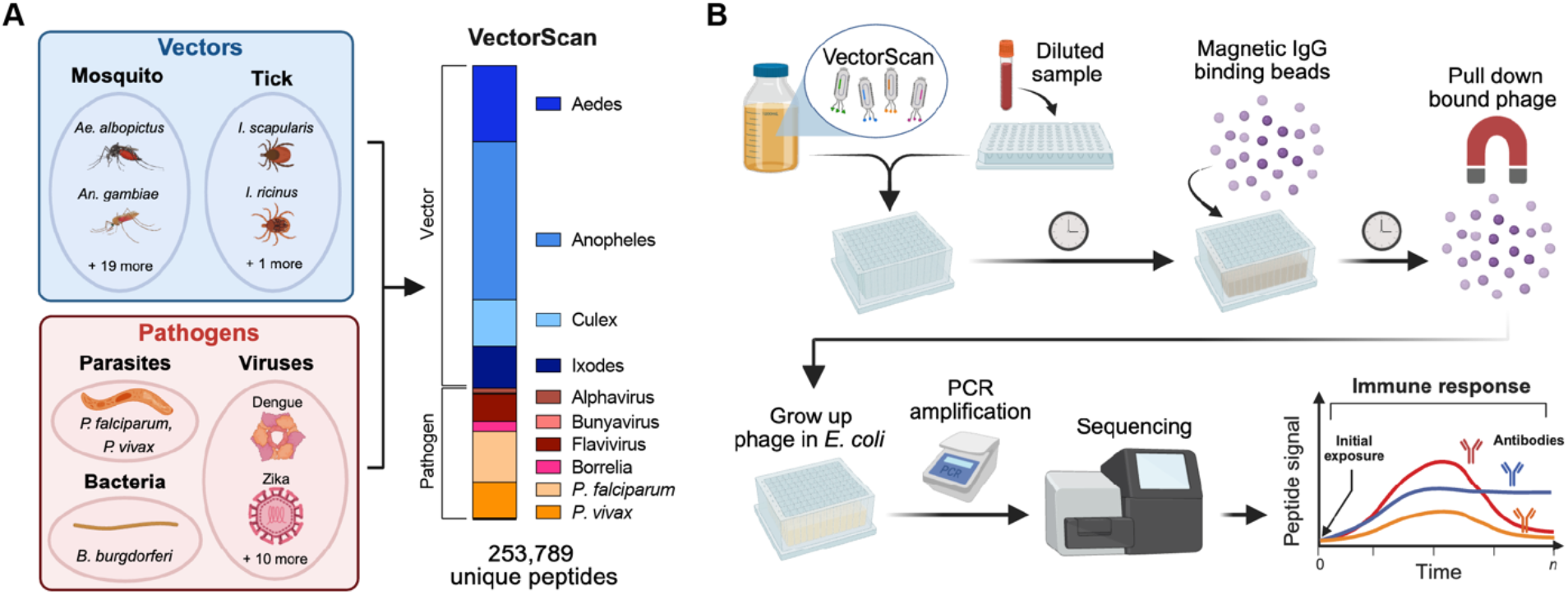
Overview of VectorScan and the PhIP-Seq pipeline. **(A)** VectorScan includes peptides from both pathogens and the sialomes of their vector carriers. This pilot library, focused on mosquito- and tick-borne illnesses, incorporates peptides from mosquitoes (*Aedes*: 42,484 peptides from 2 species, *Anopheles*: 88,271 from 17 species, *Culex*: 26,121 from 2 species), ticks (23,347 from 3 *Ixodes* species), arboviruses (Alphaviruses: 2,701 from 3 species, Bunyaviruses: 632 from 2 species, Flaviviruses: 15,205 from 7 species), bacteria (5,660 from *Borrelia burgdorferi*), parasites (28,438 from *P. falciparum* and 20,417 from *P. vivax*), and controls (412 from common human viruses and 100 containing random sequences). The fraction derived from control sequences is not labeled due to scale but is included in the final peptide count. **(B)** In Phage Immunoprecipitation Sequencing (PhIP-Seq), small peptides derived from specific antigens are encoded into bacteriophage, which express the protein on their surface. These peptides act as a library of potential linear epitopes. Once synthesized, the resulting library is packaged into phage, amplified, and subsequently incubated with samples containing the antibodies of interest (e.g., human serum). The peptides on the phage surfaces bind to antibodies targeting the antigen from which the sequence was derived. These antibody-phage complexes are then pulled down, and the peptide-containing regions of the captured phage are amplified using PCR and sequenced to generate seroreactivity profiles.

Previous studies have demonstrated that robust antibody responses to viruses can be measured using phage display. Pertinently, phage libraries have been designed to assess differential immune responses among arboviruses^13^, and flaviviruses specifically^15^. However, these libraries were composed entirely of virus-derived peptides and not vectors or non-viral VBDs, whereas arboviruses account for only 14,365 out of 249,243 peptides (5.76%) in VectorScan. To determine whether the wide breadth of the VectorScan library compromises our ability to detect viral exposure, we screened sera from three non-human primates (NHPs), each exposed to one of three distinct dengue viruses (strains of DENV1, DENV2, and DENV4) (**Fig. 2A**). The resulting VectorScan profiles (normalized sequencing counts of each peptide in the library) were unique to each NHP and longitudinally stable (**Fig. S2A**). To determine which epitopes drive humoral responses to DENV, we identified peptides that seroconverted within each NHP, meaning the peptide was seronegative at baseline but seroreactive post-exposure (see **Methods** and **Supplementary Methods**). We observed that the fraction of seroconverting DENV-derived peptides (**Fig. 2B, S2B-G**) was three to seven times higher than what would be expected at baseline in each NHP (based on Odds Ratio; p<2.2e-16 for each NHP; Fisher’s exact test; **Fig. 2B**, inset). Furthermore, despite known challenges in cross-reactivity among flaviviruses (particularly Zika virus, which shares high genetic and structural homology to DENV), this serological response enrichment was the greatest for DENV in each NHP, indicating that we could detect species-specific antibody responses.

**Figure 2:**
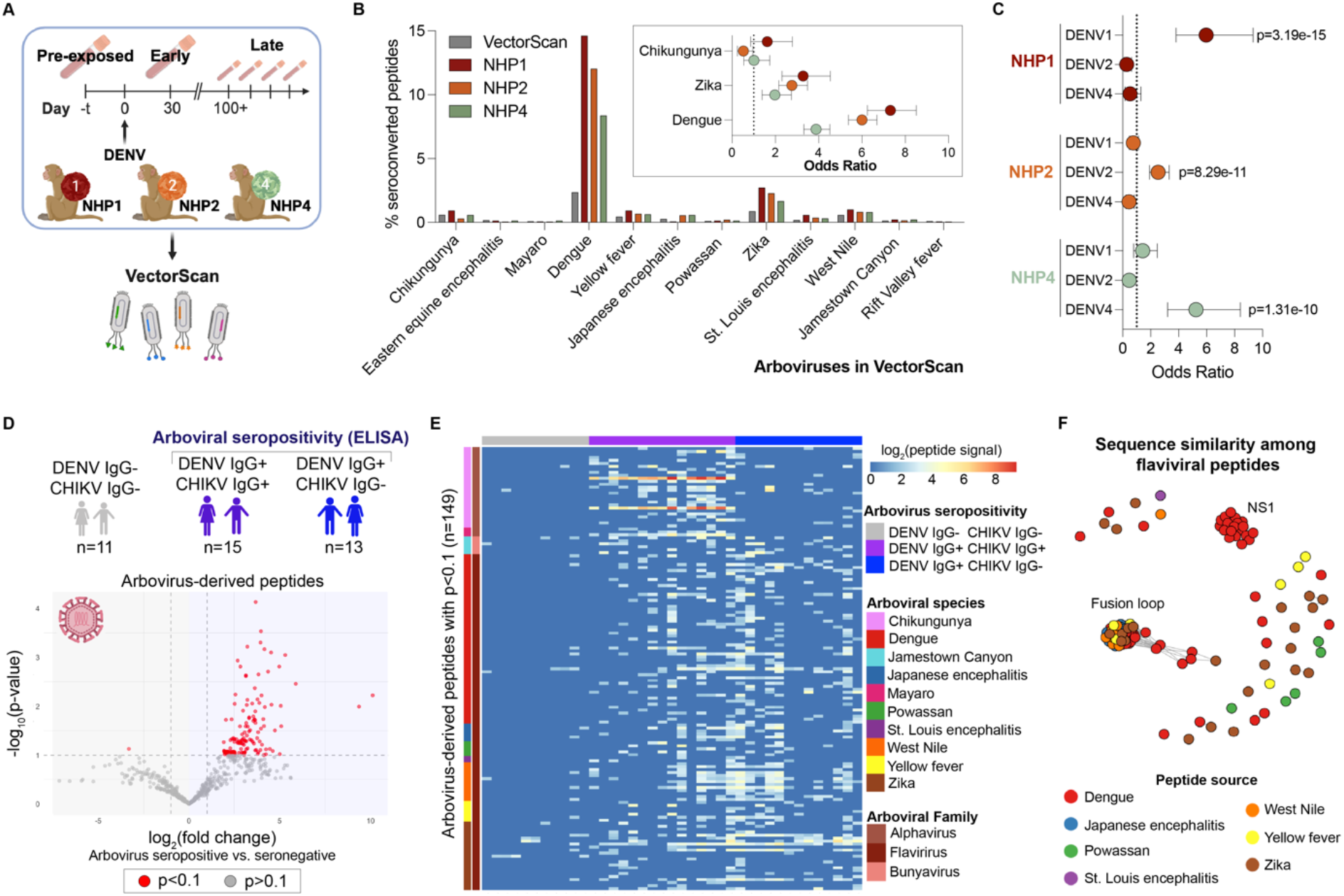
Epitope profiles in individual samples are reflective of virus-specific antibody responses and can inform serotyping. **(A)** Sera was sourced from three non-human primates (NHPs), each exposed to a different DENV strain of distinct serotypes (NHP1: infected with dengue virus 1 Nauru/West Pac/1974 [DENV1]; NHP2: infected with dengue virus 2 Thailand/NGS-C/1944 [“DENV2”]; NHP4: infected with dengue virus 4, Dominica/814669/1981 [“DENV4”]. Three samples were available per NHP: “Pre-exposed” samples collected prior to infection, “Early-immune” sera collected 30 days post-infection, and “Late-immune” sera collected more than 120 days post-infection. **(B)** The set of peptides that seroconverted post-exposure was identified within each NHP. Among these peptides, the proportion derived from dengue virus, specifically, was increased compared to the baseline percentage of representation in VectorScan. This overrepresentation of this DENV fraction was statistically significant in each NHP (p<0.0001, Fisher’s exact test). **(C)** Fisher’s exact tests were used to evaluate whether serotype-informative peptides from the infecting strain were overrepresented in each NHP. The strength and direction of the association is indicated by the Odds Ratio (OR); an OR>1 suggests overrepresentation. Error bars represent the 95% confidence interval. **(D)** Serum samples from endemic areas had pre-existing metadata of IgG seroreactivity to DENV and CHIKV. A subset of individuals who were seropositive for both viruses, seropositive for DENV only, or seronegative for both were screened with VectorScan. The signal from 626 arbovirus-derived peptides that were seroreactive in at least three people was compared between DENV IgG+ and IgG-individuals using a Student’s t-test on a peptide-by-peptide basis. Peptide signal from non-seroreactive peptides was artificially set to 0 before applying the t-tests. 149 peptides had a significant difference in group-wise means (p<0.1) between the two groups. **(E)** Peptide signal from peptides with p<0.1 were log transformed (log_2_(FC+1)) and plotted using the *pheatmap* package in R. **(F)** Sequence similarity among the flavivirus-derived peptides was quantified using *blastp* and visualized with network graphs.

After detecting anti-DENV antibody responses, we sought to determine whether regions of the proteome were immunodominant. Using *blastp*, we mapped each DENV-derived peptide in the library to the three strains used to infect the NHPs (**Fig. S2H**). Across all three NHPs, the majority of seroconverted peptides mapped to Envelope protein (E), non-structural protein 1 (NS1) and non-structural protein 3 (NS3), in agreement with previous studies (**Fig. S2I-L**).^15^ Within these highly targeted proteins, some specific regions were seroreactive in all animals regardless of the infecting viral serotype, such as E 280-365, which has homology to all three DENV strains (**Fig. S2M**) and directly precedes the pan-flavivirus fusion loop, and NS1 840-924, a region with less conservation across DENV strains that is nonetheless targeted in all NHPs. However, other regions were highly targeted in a particular NHP (e.g., E 610-645 in NHP1 and E 500-580 in NHP4), which we hypothesized was at least partially due to strain-specific humoral responses, in addition to expected individual heterogeneity. To deconvolute the basis of these inter-NHP differences, we mapped every DENV-derived peptide in VectorScan to each strain used for inoculation. After performing a threshold analysis of sequence homology (**Fig. S3**), we defined peptides as “serotype-informative” if they had at least 85% identity to one strain and less than 80% identity to the other two strains. Based on this definition, 2,668 of 5,876 DENV-derived peptides from across the DENV proteome (**Fig. S4A-D**) were serotype-informative (DENV1: n=891; DENV2: 1,192; DENV4: 585). To determine whether each NHP had serotype-informative immune responses, we identified peptides that only seroconverted in one of the NHPs (**Fig. S4E**) and tested whether serotype-informative peptides from the infecting strain were overrepresented. We observed significant representation of the infecting serotype within each NHP (**Fig. 2C, S4F**; DENV1-specific peptides in NHP1: Odds Ratio (OR)=5.97, Fisher’s exact test (FET) p=3.19e-15; DENV2-specific peptides in NHP2: OR=2.53, FET p=8.29e-15; DENV4-specific peptides in NHP4: OR=5.24, FET p=1.31e-10), suggesting that an immune response specific to the serotype that is responsible for infection is detectable. This finding indicates that the short linear epitopes in the library can effectively distinguish antibody-based infection signatures from highly similar DENV serotypes.

To evaluate the real-world ability of VectorScan to detect historical exposures to vector-borne pathogens, we screened serum collected before 2019 by the CDC Dengue Branch from individuals living in dengue endemic areas where there was no prior Zika transmission. Each sample was screened by the CDC for IgG against DENV and CHIKV via ELISA. A total of 28 individuals were seropositive for either arbovirus (IgG+) and 11 were seronegative for both (IgG-). Importantly, these individuals were not acutely ill, and there was no information about when their infections occurred. We performed Student t-tests on individual peptides for all arbovirus-derived peptides that were seroreactive in at least three people (n=626 peptides) to identify those with elevated signal (fold-change between normalized read counts for each replicate vs. no-serum controls) in the arbovirus IgG+ vs. IgG-individuals. A total of 149 peptides exhibited a significant difference between the two groups (p<0.1, **Fig. 2D**). Among these, we observed strong clustering that corresponded to chikungunya (CHIKV) IgG seropositivity, where alphavirus-derived peptides had increased signal specifically in CHIKV IgG+ individuals (**Fig. 2E**). In alignment with expectation, flavivirus-derived peptides had increased signal among the DENV IgG+ individuals, regardless of CHIKV seropositivity status (**Fig. 2E**). Notably, one of the double IgG-individuals also exhibited broad seroreactivity to the flavivirus-derived peptides that emerged from this analysis. It is possible that this sample was an ELISA false-negative, or that the individual had been infected with another cross-reactive flavivirus. We probed for common epitopes underlying these arbovirus-associated signatures as in previous PhIP-Seq studies, using *blastp* to cluster peptides based on sequence identity.^28,29^ We found that most of the alphavirus-derived peptides mapped to the CHIKV E2 glycoprotein (**Fig. S5A**), which is a known immunogenic region commonly used as the antigenic target in diagnostic assays. Among the flavivirus-derived peptides, cluster analysis revealed immunodominant signals from two regions (**Fig. 2F**). The first cluster (Cluster 1) consists of peptides from a wide range of flaviviruses that map to the flavivirus fusion loop (**Fig. S5B**). The broad cross-reactivity of the fusion loop has been explored using the previously published ArboScan library.^13^ However, interestingly, this region was more highly targeted in the humans than the NHPs, where regions just upstream of flavivirus loop were immunodominant (**Fig. S1H**). Cluster 2 is composed only of DENV-derived peptides (**Fig. S5B**) mapping to the region of NS1 that was also immunodominant among the NHPs (**Fig. S5C**).

Having demonstrated that epitope profiles could discriminate between arbovirus species, and even between highly similar DENV serotypes, we sought to investigate whether the vector-derived portion of the library could detect antibody signatures of vector exposure. We obtained samples from a clinical trial at NIH in which healthy volunteer adults living in the Washington, D.C. metropolitan area experienced four 10-minute feedings by up to ten uninfected *Anopheles gambiae* or *Aedes aegypti* mosquitoes, with each feeding approximately two weeks apart (**Fig. 3A**).^18^ We screened samples from Day 0, prior to these successive exposures, and Day 44, after all four exposures. To allow for heterogeneity in mosquito exposure among volunteers prior to the study, rather than require a peptide to be seronegative at baseline (as was done for the NHPs), we leveraged the longitudinal stability of each volunteer’s antibody repertoire to identify all peptides with increased signal between Day 0 and Day 44 (see **Supplementary Methods** for details). Within the resulting sets of “post-exposure” peptides for each person, each peptide was classified as either “seroconverted”, meaning it was seronegative at Day 0 but seroreactive at Day 44, or “boosted”, meaning it was already seroreactive at Day 0 but had increased signal at Day 44 (**Fig. 3B**). The total number of post-exposure peptides with increased reactivity varied among individuals (mean: 1,652, range: 1,240-1,981) and peptides derived from mosquitoes were significantly overrepresented in 70% (16/23) of volunteers (**Fig. 3B, Fig. S6A**, yellow shading indicates Fisher’s exact test p<0.05). We next restricted the analysis to peptides that seroconverted within each individual and found that while the number of seroconverted peptides still varied markedly among individuals (mean: 722.17, range: 487-1,151), focusing specifically on signal that emerged post-exposure resulted in significant overrepresentation of mosquito-derived peptides in a larger fraction of volunteers (20/23, **Fig. 3C, Fig. S6B**). This suggests that *de novo* immune responses elicited by repeat mosquito exposure disproportionately recognized epitopes within mosquito-derived peptides relative to non-mosquito peptides in the library, enabling evaluation of recent acute exposure at an individual level. Interestingly, across all volunteers, the difference in signal between Day 0 and Day 44 is significantly higher among mosquito-derived peptides that exhibit boosting compared to those that seroconverted (FDR-corrected p_adj_<1e-05 for each volunteer, Wilcoxon rank-sum test; **Fig. 3D**), which may indicate that previously encountered epitopes elicit a more robust antibody response to acute exposure driven by immunological memory. Furthermore, while reactivity to the same boosted peptides was more frequently observed in multiple individuals (**Fig. 3E**), 79.2% of all peptides with increased reactivity and 91.5% of seroconverted peptides, specifically, were only observed within a single individual (**Fig. 3E**), highlighting the variability of individual human responses to a common acute stimulus.

**Figure 3:**
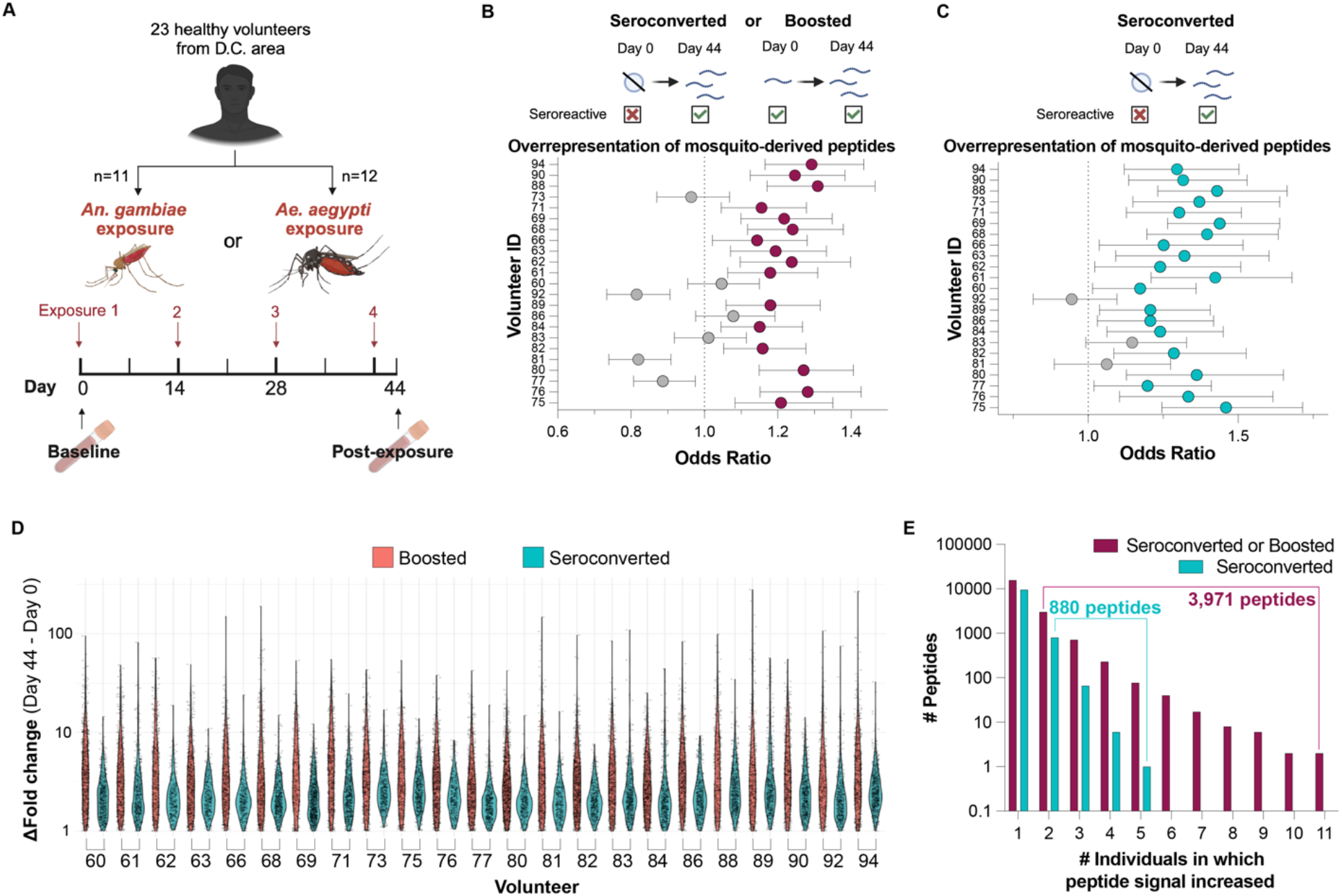
Increased recognition of mosquito-derived epitopes after controlled vector exposures. **(A)** Healthy adult volunteers underwent four feedings by uninfected *An. gambiae* or *Ae. aegypti* mosquitoes. Plasma samples from Day 0, prior to these successive exposures, and Day 44, after all four exposures, were screened with VectorScan. **(B)** Peptides with an increase in signal (fold change values between replicates and PBS controls) were identified within individuals. Peptides were categorized as “boosted” if they were seroreactive at Day 0 and “seroconverted” if they were seronegative at Day 0. Fisher’s exact tests were performed to evaluate whether mosquito-derived peptides were overrepresented among these peptides. The strength and direction of the association are indicated by the Odds Ratio (OR); an OR>1 suggests that the mosquito-derived peptides are overrepresented. Purple dots indicate that the association is significant (p<0.05). In **(C)**, the set of peptides being analyzed was limited to those that seroconverted between Day 0 and Day 44. Teal dots indicate that the association is significant (p<0.05). Error bars in (B) and (C) represent the 95% confidence interval. **(D)** The difference in signal (difference in fold change values) was determined for each peptide contributing to the signals in panel B, stratifying between peptides that were boosted versus seroreactive (but that nonetheless had increased signal post-exposures). **(E)** Quantification of the number of volunteers in which these peptides had increased signal.

To better understand the nature of the antibody response to acute mosquito exposure, we explored the functional profile of the proteins from which these post-exposure peptides were derived. We found that 3,971 peptides had increased post-exposure signal in at least two individuals (**Fig. 3E**). Three of the most highly represented protein classes of these peptides were serine proteases (IPR001254, n=312 peptides), proteins with chitin-binding domains (IPR002557, n=249), and proteins with Immunoglobulin-like domains (IPR036179, n=162; **Fig. 4A**). Although serine protease-derived peptides were a large fraction of the signature, serine proteases, as a group, were not overrepresented compared to the overall library (*i*.*e*., for IPR001254, p=1.39e-06, OR=0.76, Fisher’s exact test). Rather, peptides with chitin-binding domains, from insect cuticle proteins, and immunoglobin-like proteins were significantly overrepresented (IPR036508: p=0.0003, OR=1.27, IPR000618: p<0.0001, OR=2.98, IPR036179: p=0.016, OR=1.19, Fisher’s exact test; **Fig. 4B**). Recent studies have identified cuticle proteins as insect- and arthropod-derived allergens, but their ability to elicit IgG responses is not well-characterized.^30–32^ A previous study that used a cDNA library derived from house dust mites (an arthropod) to identify proteins with IgE-reactivity also found that chitin-binding domain containing proteins and chitinases (which belong to glycoside hydrolase family 18 [IPR001223], another overrepresented group) were allergens.^33^ This suggests that the protein families that emerged in these analyses may represent elements of the humoral host immune response to mosquitoes that have been understudied with regard to their broader immunogenicity and have value as targets for future work.

**Figure 4:**
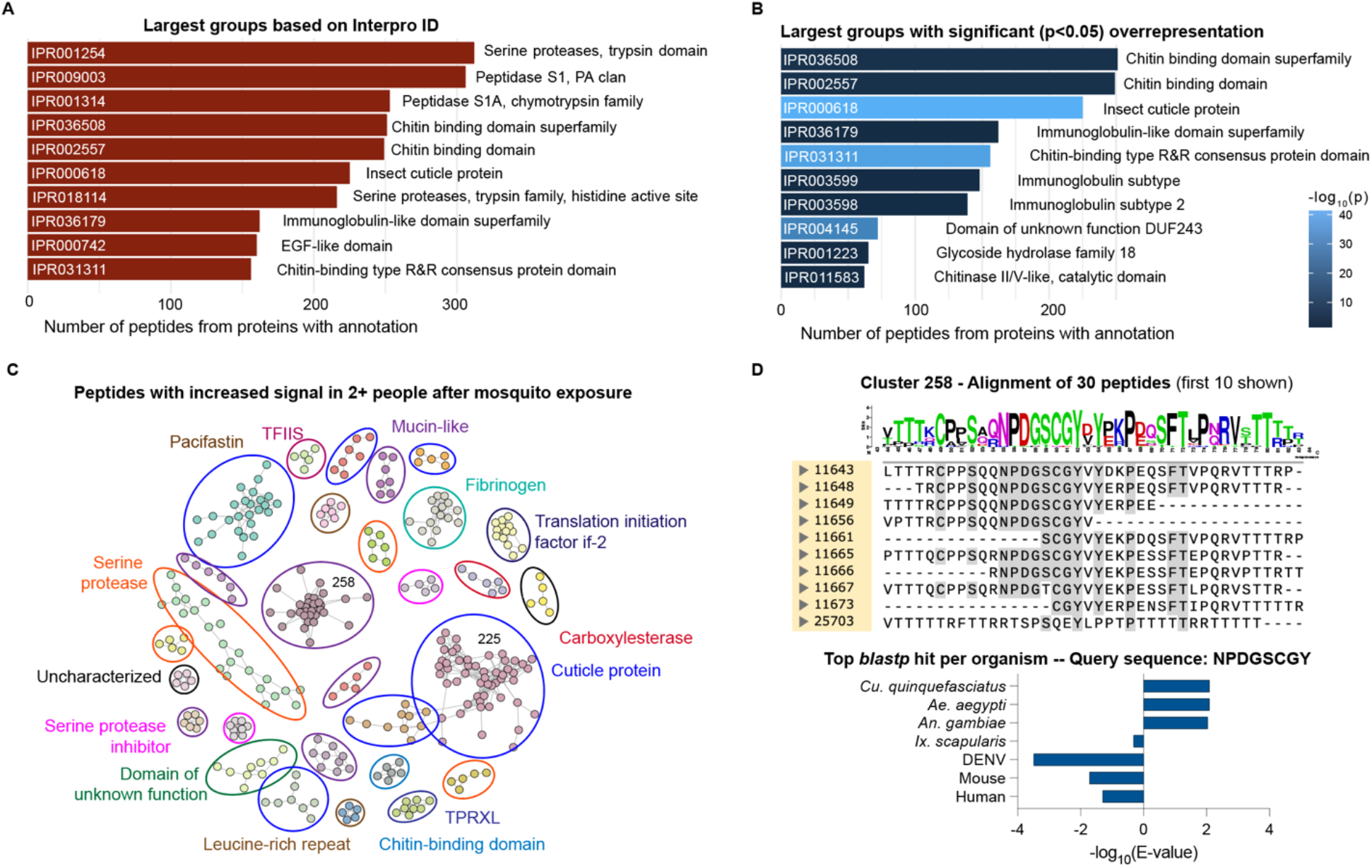
A wide range of protein classes and distinct epitopes are enriched post-mosquito exposure, suggesting a multifactorial antibody response. The frequency of InterPro annotations at the protein level **(A)** and the enrichment of these domains relative to the entire mosquito-derived portion of VectorScan **(B)** were analyzed for these 3,971 peptides in the post-exposure signature. In **(A)** the bars are ranked in order of size, while **(B)** is ranked by descending -log_10_(p). The top ten groups are shown in each panel. Annotations were not condensed based on hierarchy within InterPro. **(C)** Immunogenic epitopes were identified by identifying peptides with conserved sequences via *blastp*. Network graph analysis was used to cluster peptides with high similarity. Clusters with at least five members were visualized. Manual protein-level annotations for clusters are labeled. **(D)** Peptide sequences from cluster 258, which are from mucin-like proteins, were aligned with MUSCLE in SnapGene. A logo plot was generated based on the alignment, and amino acids with at least 70% conservation across all sequences are highlighted in gray. The longest segment of the consensus was used as a *blastp* query against several target organisms in NCBI. The log_10_(E-value) of the top hit is plotted; higher values correspond to a lower E-value and therefore represent a more significant association between the query sequence and the organism of interest.

To probe these candidate proteins more deeply, we performed a detailed analysis of epitope sequences within these proteins to identify immunodominant epitopes. In PhIP-Seq studies, the enrichment of multiple peptides containing similar motifs increases the confidence that the re-occurring sequence is an immunogenic epitope.^34–36^ We used *blastp* to identify Day 44 enriched peptides with high similarity to at least one other hit (n=1,388, 35.0% of the original peptide set), organized them into clusters (those with five or more members are visualized in **Fig. 4C**; all cluster compositions are available in **Table S3**), and visualized them with network graphs, allowing us to identify targets of the host antibody response. Many distinct clusters emerged from this analysis, indicating that a wide breadth of epitopes elicit immune responses. Clusters of peptides from cuticular proteins were particularly interesting, because the cuticle is a fundamental structural component of the exoskeleton that is unique to arthropods. The largest of these clusters (#225), contains peptides that are valine/proline rich with similarity to proteins in the cuticular protein of low complexity (CPLCP) family.^38^ Another large cluster, Cluster 258, contains peptides from proteins that are annotated as mucin-like (**Fig. 4C**) and contain a consensus sequence, “NPDGSCGY” (**Fig. 4D**). The top *blastp* hits across the entire NCBI database for this sequence are from mosquitoes (comparisons among selected model organisms are shown in **Fig. 4D**) and appears to be conserved across multiple mosquito genera, as indicated by similar E-values and query coverage, suggesting it could be a cross-reactive epitope. Consensus sequences such as these could be incorporated into future multiplexed panels or ELISAs to broadly assess recent mosquito exposure at an individual level, or leveraged in anti-vector vaccine strategies.

Finally, we sought to validate these findings in the setting of natural exposure. Because an arboviral infection typically necessitates prior mosquito exposure, we explored whether there were measurable differences in antibody responses to mosquito-derived epitopes between CDC samples which were IgG+ for either DENV or CHIKV, or IgG-for both. We performed Student t-tests for all mosquito-derived peptides that were seroreactive in at least one person (n=29,736 peptides) and observed higher enrichment of peptide signal in the IgG+ group (**Fig. 5A**). Among the 902 peptides that were associated with exposure (p<0.1), the difference in signal appeared to be driven by high peptide seroreactivity in one group and a lack of seroreactivity in the other, rather than a difference in the magnitude of signal among broadly seroreactive peptides (**Fig. 5B**). These group-wise differences are bi-directional; while most peptides have higher seroreactivity among the IgG+ individuals, a small subset of mosquito-derived peptides have increased seroprevalence/seroreactivity among individuals who were seronegative for both arboviruses. Because CHIKV and DENV are transmitted by *Aedes* mosquitoes, we explored whether *Aedes*-specific responses were enriched within this set of peptides. Among the 50 peptides with the smallest p-values, the proportion of *Aedes*-derived peptides was significantly overrepresented (p=0.0064, Fisher’s exact test) relative to the full exposure-associated signature (902 peptides), but *Anopheles*- and *Culex*-derived peptides were not (**Fig. 5C**). However, when we compared the origin of the exposure-associated signature to the original set of peptides used to identify this set (n=29,736), this genus-level enrichment fades. The epitope sequences underlying the exposure-associated signature revealed that the largest cluster (cluster 64) contained the same “NPDGXCGY” motif as cluster 258 (which mapped to mucin-like proteins) from the NIH cohort (**Fig. 5D,E**), though the exact peptides differed between the respective clusters. To directly evaluate the commonalities between the epitopes that both clustered and exhibited immunogenicity in at least two people after acute mosquito exposure in the NIH cohort (n=1,388 peptides), with those included in the CDC samples that clustered and were associated with a known historical arbovirus exposure (n=129 peptides), we re-performed this network graphing analysis (using all 1,517 peptides). Visualization revealed additional clusters containing epitopes from both cohorts (circled, **Fig. 5F**), representing other shared antigenic targets between the two cohorts. Aside from mucins, some of these epitopes were derived from cuticle proteins, serine proteases, and proteins with chitin-binding domains.

**Figure 5:**
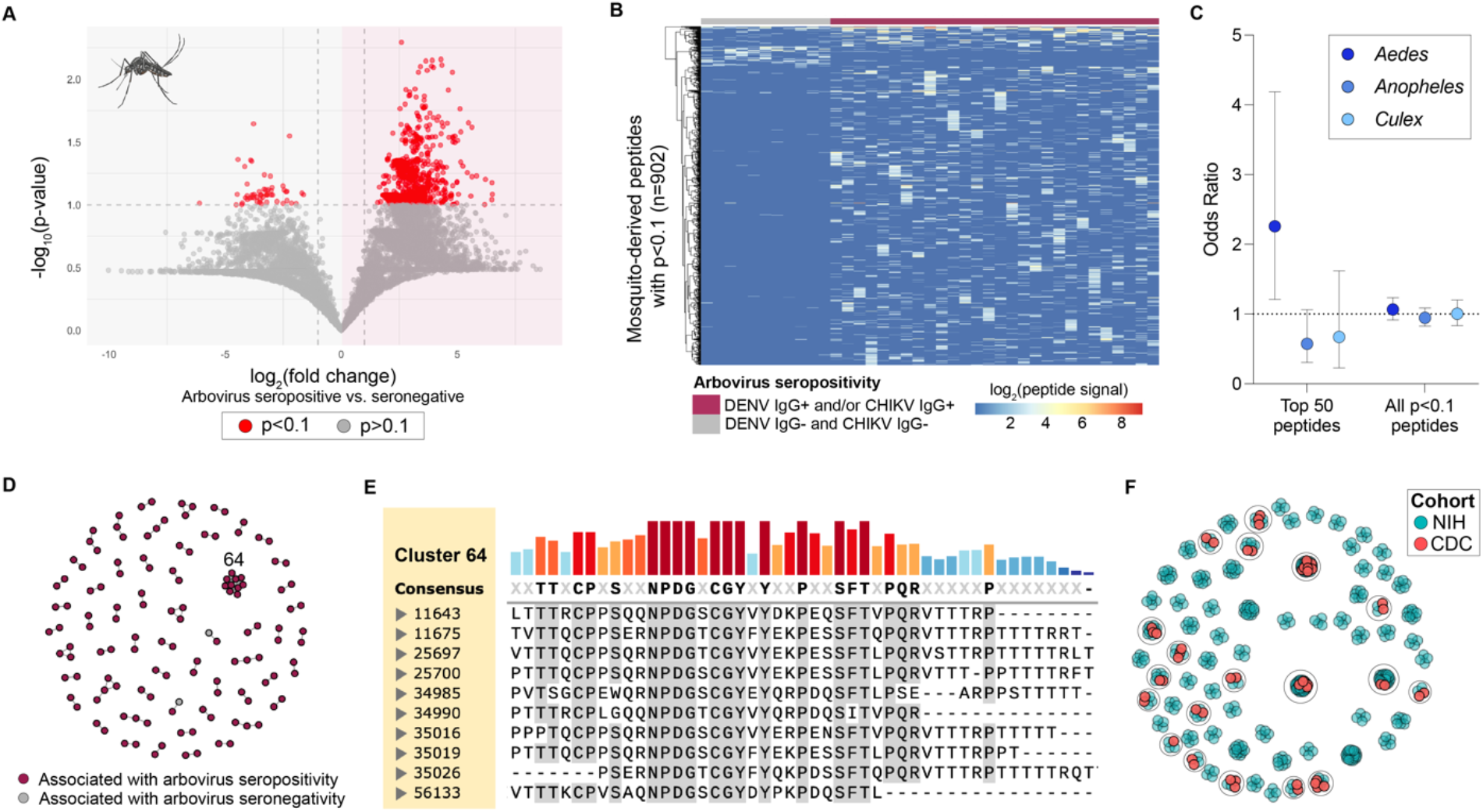
A subset of mosquito-derived epitopes has differential signal among individuals with ELISA seropositivity for a historical arbovirus infection versus those who are seronegative. **(A)** The peptide signals (fold change of normalized counts from each sample over mock-IP controls) of 29,736 mosquito-derived peptides that were seroreactive in at least 1 person from the CDC samples were compared among individuals who were IgG+ via ELISA for CHIKV and/or DENV, or IgG-for both. The log_2_(fold change) of the average peptide signal across all individuals from each group versus the p-value of the Student t-test comparing those means was graphed for each peptide. Peptide signal from non-seroreactive peptides was artificially set to 0 before applying the t-tests. Each point represents a peptide; red indicates a p<0.1 and a |log_2_(fold change)|>1, which is considered significant. **(B)** A heatmap was created based on the signal of each peptide that reaches this significance threshold. Each column is a different individual. **(C)** Fisher’s exact tests were performed to calculate the overrepresentation of each mosquito-family among the 50 peptides with the smallest p-values versus the 902 peptides that achieved significance. A second set of tests were performed to compare the mosquito-family representation among these 902 peptides versus the original set of 29,736 peptides that were seroreactive in at least one person. Error bars represent the 95% confidence interval. **(D)** Clustering analysis of the significant peptides identified sets of peptides with similar sequences. **(E)** The sequences from the largest cluster (#64) were aligned via MUSCLE in SnapGene. Gray highlighting indicates at least 70% consensus among the aligned peptides. **(F)** The peptides used to create the network graphs in Fig. 4C and Fig. 5D were combined and re-clustered to evaluate the degree of similarity between the immunogenic epitopes that arose from the controlled mosquito exposure and natural exposure cohorts. Each node is a peptide; blue nodes are peptides from the NIH cohort (where there was controlled mosquito exposure) and red nodes are from the CDC samples (where there was presumed natural exposure based on arbovirus IgG seropositivity). Clusters with at least four members were visualized.

## Discussion

We have designed a custom phage display library, VectorScan, to query antibody repertoires against a wide range of organisms responsible for both the pathogenesis and transmission of vector-borne diseases. Whereas classic serological tools such as ELISAs and lateral flow assays often fail to distinguish between closely related flaviviruses due to cross-reactivity,^39^ we have added to the growing body of work that demonstrates how broad interrogation of arboviral proteomes with highly multiplexed assays can discriminate among flavivirus species as well as DENV serotypes.^13,15,40^ We also demonstrated that the seroreactive peptides we detected have translational potential by screening samples from individuals in endemic areas who had natural arbovirus exposures, and observed reactivity to epitopes from CHIKV and DENV that aligned with ELISA-based diagnoses of prior arbovirus infections. Leveraging the unique duality of VectorScan, we also observed that these field samples had significantly different mosquito-derived epitope profiles from those who were seronegative for CHIKV and DENV. Notably, the hits with the most significant differences between these groups were overrepresented by *Aedes*-derived peptides, suggesting greater past vector exposure in the arbovirus seropositive individuals. Furthermore, some of the epitopes associated with known arbovirus exposure were found to be immunogenic among individuals with known, acute, controlled mosquito exposures. Overall, we have demonstrated the utility of a tool to simultaneously profile anti-vector and anti-pathogen antibody profiles.

Importantly, the wide breadth of pathogen and vector proteomic space we were able to simultaneously profile with VectorScan does not compromise the specificity of the inferred exposure profiles relative to more focal libraries and suggests that further expansion and refinement of the library will be practical. For example, the patterns of antibody reactivity to DENV-derived peptides across the proteome that we observed in the infected NHPs were remarkably consistent to those generated by an earlier study that used a flavivirus-specific phage library.^15^ These patterns were also reproducible at the epitope-level; the arbovirus-focused ArboScan library was used to identify epitopes that were associated with DENV neutralization, such as an epitope within the envelope protein (“KGVTQNGRLITANP”) which was correlated with DENV1 neutralization, specifically.^13^ In our study, we also observed that the density of post-exposure peptides containing this epitope was highest in the DENV1-infected NHP, and that many of the peptides driving this signal were serotype-informative (meaning they had higher sequence homology to DENV1 than the other serotypes). Furthermore, among our DENV-exposed human samples, we detected high cross-reactivity of flavivirus-derived peptides mapping to the fusion loop, which was also observed using ArboScan, further supporting our ability to reliably query anti-arbovirus responses despite the relatively small representation of these organisms in the overall VectorScan library.

By dedicating most of the antigenic space in the library to vector-derived peptides, we created serological profiles against a very broad array of vector-derived peptides, allowing us to provide additional evidence for the extreme inter-individual heterogeneity of the antibody response to arthropod vector exposure. As was reported in a recent study employing a yeast display library of tick proteins,^17^ we observed high inter-individual variation following controlled exposure to bites from two mosquito species, even though each volunteer had identical exposure timelines to a known mosquito species. This individual heterogeneity has been a challenge for the use of single-peptide mosquito-derived antigens as exposure biomarkers at an individual level,^8,41^ and underscores the value of screening large panels of antigens in developing effective vector-based surveillance tools and potentially anti-vector vaccines. In addition, both a recently published tick study^17^ and this work demonstrate that an unexpectedly wide range of the arthropod proteome is immunogenic in humans exposed to vector bites. We found that cuticular proteins, structural proteins, and other non-canonical salivary proteins, which are historically overlooked in vector-based vaccines and surveillance tools, were immunogenic. It is possible that cuticular and other structural proteins are being introduced into the human host via shedding from the proboscis during probing, or that mucins are being co-delivered through saliva after being transported from the midgut to the salivary glands. Regardless of the mechanism, such proteins may be valuable targets for further study in pursuit of vaccine targets that could disrupt transmission of VBDs. In particular, the mosquito-derived epitopes that were found to be associated with arbovirus infection history may serve as promising starting points.

While we took care to include peptides from proteins traditionally used as serosurveillance biomarkers of mosquito exposure, such as AeD7L1^7^ and salivary gland 6^8^, we did not observe significant changes in seroreactivity among these proteins within the individuals in the NIH cohort. We suspect that this is due to multiple factors. First, to participate in the clinical study, individuals needed to have baseline negativity to salivary gland homogenate (SGH) via ELISA, and it has been shown that seroreactivity to SGH correlates with biomarkers such as AeD7L1. Then, each person only underwent four exposure sessions, which is a low level of exposure compared to settings where serosurveillance via salivary biomarkers is the most successful. For example, it was recently shown that two controlled exposures to *Ae. aegypti* were unable to elicit IgG responses against AeD7L1 and AeD7L2 in individuals from New York state.^42^ However, IgG levels against these two biomarkers were significantly higher in individuals from Thailand than the New York residents. The need for high levels of exposure to mosquitoes to elicit meaningful immune responses to these proteins may explain their lack of prominence among our results. However, seroreactivity to such biomarkers was also low among the CDC samples, which were collected in areas with endemic arbovirus transmission. It is possible that the lack of seroreactivity is due to the inability of PhIP-Seq to effectively detect conformational epitopes. In addition, it may be that immunogenicity to these oft used markers varies due to age, ethnic background, geography, and other demographic factors, which are often controlled for on a study-by-study basis, making them more suitable for serosurveillance in specific populations. By broadening the set of proteins used to quantify exposure histories via techniques such as PhIP-Seq or other highly multiplexed assays, a wider range of biomarkers may be identified.

Another limitation of this work is the relatively small sample size, particularly for the DENV-exposed NHPs, for which we only had samples from three animals. PhIP-Seq studies benefit from large sample cohorts with known exposure histories to strengthen confidence in disease-associated hits, and the importance of including healthy control cohorts with 50 or more individuals has been evaluated.^43^ However, such samples, especially with known exposure histories, are challenging to collect and we therefore focused here on longitudinal samples with known vector or virus exposures, which allowed us to customize our analytical pipeline to take full advantage of the longitudinal stability of PhIP-Seq repertoires and identify peptides with increased signal only after exposure to a given organism. Furthermore, to demonstrate feasibility in non-longitudinal samples, we performed initial testing of field samples with known arbovirus exposure, which also gives us confidence in the library performance. However, additional work will be needed to further explore the utility of VectorScan as a diagnostic or serosurveillance tool, including screening samples from larger patient cohorts, and including samples from people or animals exposed to malaria, *Borrelia*, and other VBD-relevant organisms, to validate the other portions of the library.

The existing VectorScan library has the capacity to simultaneously survey antibody responses to both vectors and vector-borne pathogens, a technological advantage over existing serological assays. For example, the ability to link vector exposure to disease prevalence can be used to understand how shifts in vector population density and range, due to climate change or vector interventions, influence the risk of disease outbreaks by generating individual- and population-level seroprevalence data to inform epidemiological and ecological models. Finally, the multiplex capabilities afforded by phage display dramatically scale up the information yielded by a single sample, supporting vaccine candidate and biomarker discovery, and enabling the creation of biomarker panels that accommodate highly heterogeneous individual immune responses.

## Resource availability

### Lead Contact

Requests for additional information should be made to Dr. Daniel Neafsey.

### Materials Availability

- This version of the VectorScan library was constructed using a T7 backbone proprietary to the Elledge Lab at Harvard Medical School, which was shared via a material transfer agreement. Therefore, we cannot share the library without consent from Stephen Elledge, though reasonable requests will be considered and handled appropriately.

### Data and Code Availability

- Data: All data (e.g., de-multiplexed peptide counts for each sample) are available on the Harvard Dataverse (https://doi.org/10.7910/DVN/IBY5BY). The raw fastq files are available through NCBI (PRJNA1306026).
- Code: All code used to conduct analyses can be found on GitHub (https://github.com/Fortune-Lab/VectorScan-Manuscript-1).

## Author contributions

M.A., S.M.F., and D.N., conceptualized the project. M.A., S.M.F., and D.N. planned experiments and drafted the manuscript, incorporating feedback from all authors. M.A., D.S., and E.P. performed all wet lab experiments. M.A. and A.C. created the peptide library. M.A. analyzed the data. J.R.L., J.V., M.J., F.O., L.W., and S.K. collected and provided human samples on behalf of the NIH and CDC, and provided feedback on data analysis.

## Acknowledgements

Funding for this work was supported by generous gifts from Lisa and Mark Schwartz, Rick Salas and John Kang. This work was also supported in part by the National Institutes of Health (NIH), Office of Clinical Research, Bench to Bedside Grant and was supported in part by the Intramural Research Program of the NIH for J.V., F.O., and L.W. The contributions of the NIH author(s) were made as part of their official duties as NIH federal employees, are in compliance with agency policy requirements, and are considered Works of the United States Government. However, the findings and conclusions presented in this paper are those of the author(s) and do not necessarily reflect the views of the NIH or the U.S. Department of Health and Human Services. J.R.L. was supported by the NIAID Transition Program in Clinical Research. Samples from dengue endemic areas were generously provided by the United States Center for Disease Control and Prevention Dengue Branch, with special thanks to Freddy Medina, Maile Thayer, and the rest of their team. Matthew Memoli led the NIH clinical trial that provided the mosquito-exposed samples used in this work. M.A. utilized BioRender to create elements of Fig. 1, Fig. 2A, D, and Fig. 3A-C. The T7 backbone was provided by Stephen Elledge at Harvard Medical School, and protocol assistance was provided by Mamie Li. The authors would also like to thank Ana Pinharanda, Aditi Saxena, Douaa Mugahid, and Mike Chao for their review of the manuscript.

