## Supplementary Information for "Comprehensive evaluation of antibody responses to mosquitoes and mosquito-borne arboviruses using highly multiplexed serology"

**Supplementary Methods**

All code used to generate and analyze data throughout the paper can be found in the R notebook files and GraphPad Prism 10 files that are available with this publication. Visualization was performed in R and GraphPad Prism 10.

**Design of the oligo library for the non-vector and non-viral portions of VectorScan**

Vectors: Because many vector species are non-model organisms, we selected those with well-defined sialomes to identify likely sialome proteins in other relevant but less studied species. First, we curated a set of "seed" sialomes from previous publications representing different genera of vectors that are relevant to human disease consisting of *Aedes aegypti*,[1] *Anopheles gambiae*,[2] *Culex quinquefasciatus*,[3] *Ixodes scapularis*,[4] and *Xenopsylla cheopis*.[5] Next, we curated a set of "test" datasets by downloading available genome, transcriptome, or Uniprot sequences (based on sequences available in October 2022) for various *Anopheles, Aedes, Culex, and Ixodes* species (see **Table S1** for details). We used UBLAST to find orthologs of all seed sequences in the test datasets with an e-value threshold of 1e-10 and removed duplicate sequences. We also filtered out sequences from *X. cheopis* from the list of candidates, as we used it to find orthologs of salivary proteins in the test species but focused this pilot library on ticks and mosquitoes only. We removed possible paralogs or redundant sequences by using *cd-hit* to cluster all sequences at 98% identity, resulting in 209,862 proteins. Due to the size of this initial list, we clustered progressively at 75% and 70% identity to reduce the candidate pool to 65,921 proteins. We then ran each protein through SignalP and filtered for those with a signal peptide, further reducing the pool to 11,839 protein sequences. Sequences were listed alphabetically within each FASTA, resulting in increased representation post-clustering and post-tiling for proteins from *Ae. aegypti, Anopheles albimanus,* and *C. quinquefasciatus*, which were the first species represented within each genus (**Table S1**).

Bacteria: Full protein sequences were obtained from Uniprot for *Borrelia burgdorferi* (taxonomy ID: 139), resulting in a pool of 9,251 proteins. Two strategies were then used to narrow down the pool of candidate proteins. In the first strategy, *cd-hit* was used to cluster proteins with greater than 98% identity. The resulting 3,350 proteins were input into SignalP 6.0 and the 544 proteins predicted to be secreted were selected for downstream analysis.[6] In parallel, the genes in the Uniprot list were filtered based on gene ontology for various annotations (outer membrane proteins: GO:0019867, GO:0009279; extracellular region: GO:0005576, flagella: GO:0009288). The resulting 4,143 proteins were then clustered with *cd-hit* based on 98% identity, resulting in 1,097 proteins. The protein lists from these two strategies were combined, and identical sequences were clustered using *cd-hit* to create the final set of Lyme proteins. The *pepsyn* package was then used to create tiles using the same methods as described for the viral sequences.

Plasmodium: Because Plasmodium species have large proteomes that would necessitate the creation of species-specific phage display libraries to represent the full protein space, more stringent protein selection methods were applied to target predicted immunogenic proteins. Previously, Rhaghavan et al. created a PhIP-Seq library (Falciparome) specifically for *Plasmodium falciparum,* which they screened against samples collected from individuals in the United States and Uganda.[7] Trusting the rigor of their approach, we used PlasmoDB to retrieve the sequences of all genes that yielded seroreactive peptides in their study, and mapped them back to the Uniprot, resulting in 2,939 proteins. The gene name for each entry was extracted and used to query all Uniprot entries from *P. falciparum* (taxonomy ID: 5833) and *P. vivax* (taxonomy ID: 5855). The resulting proteins from both species were combined, yielding 48,499 sequences, and *cd-hit* was used to cluster based on 95% identity to 48,390 proteins. The list was then input into SignalP, which enabled extreme downselection to 2,944 proteins, which were then tiled through with *pepsyn* as described above, yielding 40,339 peptides. To ensure that all seropositive peptides from the Falciparome screen were included in the library, we then clustered our list with the seropositive peptides using a threshold of 75% and included all Falciparome peptides that did not cluster with one of our existing sequences, resulting in a final list of 48,855 Plasmodium peptides. Notably, the sequences were used directly from the Falciparome, and each is 62-amino acids long. There are also 27 peptides with sequences derived from *Anopheles*, but these were not used for downstream analysis of the NIH cohort in this study and are included in the count for the Plasmodium/parasite/pathogen section of the library.

Arboviruses: Protein sequences from all 51,136 Uniprot entries for the viruses of interest (see **Table S1**) were compiled into a single FASTA file and *cd-hi*t was used to cluster sequences with 98% identity, distilling this list to 2,106 proteins.

Controls: To create negative control sequences, we generated 100 random 56-mers. For positive controls, peptides from previous phage libraries with high seroprevalence were selected and tiled.[8,9] To ensure comprehensive coverage of these sequences, the full proteins from which these seroprevalent peptides originated were also included.

After protein down-selection, we then created 56-amino acid peptide tiles with 28 residues of overlap using the *pepsyn* package.[10] We used *cd-hit* to reduce amino acid level redundancy of tiled peptides within each major taxonomic group based on a threshold of 95% identity.

**Construction of the VectorScan phage library**

The resulting oligo pool was synthesized by Twist Biosciences and amplified via PCR using the KAPA HiFi HotStart PCR kit (12 cycles).

- Forward primer: 5' GTCCTTAGCAGTCAATGATACGGCGTGAATTCGGAGCGG 3'
- Reverse primer: 5' GGTTACACTCCATCAAGCAGAAGACGTCTCGAGTGCAGTG 3'

The amplified library digested with XhoI and EcoRI-HF at 37°C for 60 minutes. A post-digest cleanup was performed using the Monarch PCR & DNA Cleanup Kit.

The T7F bacteriophage was provided by Steven Elledge at Harvard Medical School. It was amplified in BLT5403 and digested with SalI-HF and EcoRI-HF. The digest was cleaned up using the Monarch PCR & DNA Cleanup Kit and run on a 2% LMP gel. The band containing the T7F DNA was extracted and eluted using the Zymoclean Large Fragment DNA recovery kit. The resulting DNA was concentrated using Amicon spin filters and the concentration was measured via Qubit. The digested vector DNA and insert DNA were ligated using T4 DNA ligase at 16°C overnight in a thermocycler. The ligated product was packaged using the T7 Select Packaging Extract kit, and qPCR was used to confirm that the ligation was successful. The resulting phage library then underwent two rounds of amplification on plates. The resulting lysate was titered with a plaque assay and subsequently used for PhIP-Seq.

**Phage Immunoprecipitation Sequencing (PhIP-Seq) procedure**

We screened each sample with VectorScan in duplicate alongside no-serum controls, where the biological sample was replaced with an equal volume of PBS. First, each relevant well of a 1.1 mL 96-well plate was filled with 1 mL of TBST + 3% BSA and rotated overnight at 4°C. The following day, the phage library was prepared by combining the expanded library (~5e10 pfu/mL) with 50 ug/mL of chloramphenicol and 50 ug/mL of kanamycin. The blocking solution was removed from the plate and 1 mL of the prepared library and 10 uL of diluted sample (~2 ug IgG) was added to each relevant well and rotated overnight at 4°C. Next, immunoprecipitation was performed by adding 40 uL of a 1:1 mixture of Protein A and Protein G Dynabeads (Thermo Fisher Scientific) to each well and incubating with rotation for 1 hour at 4°C. The beads were washed 5 times with cold RIPA buffer, resuspended in M9LB+Carbenicillin in a 2 mL deep-well plate, and added to 1 mL of E. coli (BLT 5403) grown to an OD600 ~0.5. The plate was then incubated in an orbital shaker at 37°C until the wells cleared. NaCl was added to each well and the plate was centrifuged at 3,000 x g for 1 hour at 4°C. Two uL of lysate was used as input for a PCR reaction to amplify the peptide-encoding inserts. A subsequent PCR2 reaction was used to apply indices to each sample. The indexed samples were pooled and submitted for sequencing, which was performed by the Harvard University Bauer Core Facility. The adapters were trimmed from each read using *cutadapt* and aligned to the DNA sequences in the VectorScan library using *bowtie-2* (options: --very-sensitive -U) to obtain counts for subsequent analyses.[11,12]

**Determination of seroreactivity thresholds**

As described in the main text, after sequencing, normalization was performed. For Plates 14, 15 and 22 (which cover all human samples), samples were excluded from analysis if the raw read counts were poorly correlated or if one of the replicates for the sample did not reach an adequate number of reads. These sample exclusion criteria were not used for Plate 13 (NHPs) due to the small sample size. Peptide counts were first normalized by sequencing depth within each replicate (based on counts per 100,000 reads, RPK). We then calculated the fold change between the RPK of each sample (treating replicates separately) over the median RPK across the no-serum controls on a per-peptide basis (1 was added to both the numerator and denominator to prevent division by 0). The resulting normalized fold change values were used for the remainder of analyses, since they reflect both the relative abundance of each peptide within a sample and the relative number of counts above what would be expected by nonspecific binding. Consistency between normalized replicates was assessed by evaluating the Pearson coefficient between pairs. For most samples, the correlation between replicates was high (>0.8), and correlations were particularly strong for the human samples (**Fig. S6**). However, some of the NHP samples had lower Pearson coefficients, which led us to create conservative criteria for determining seroreactivity that relied on consistent signal from both replicates.

Methods for analyzing PhIP-Seq data differ, on a per-lab and even per-study basis. However, in previous PhIP-Seq studies, seroreactivity (often used interchangeably with "seropositivity") is usually defined based on exceeding a certain threshold of counts, z-score, fold change over mock-IP wells, or a combination of multiple criteria. These thresholds are adjusted on a per-study basis, which makes it difficult to assess what this threshold should be when using a new library. To address this concern, we designed the VectorScan library to contain 100 peptides with randomly generated peptide sequences. This portion of the library was included as a collective negative control, as the reactivity of these peptides will be measured in each replicate that is screened, which would allow us to normalize the VectorScan signal on a per-sample basis. We averaged the fold change counts for each peptide over both replicates from a sample and evaluated the distribution of values from each sample (**Fig. S6**). While these peptides were not rationally selected to interrogate biological signal, the exposure history of each person is vast, and we found that a small number of peptides had fold change values that were "high" relative to the background distribution of all 100 random peptides. We assume that some signal from these random peptides may represent true antibody reactivity, therefore we set a threshold for noise at 80% of the signal across all 100 peptides in each sample (**Fig. S6**; each random peptide is represented by black dots, while the 80% threshold is represented by a red dot). Essentially, if a peptide has a fold change value below this 80% threshold, it is safe to say that the signal does not represent robust biological antibody reactivity. To call a peptide "seroreactive", we took a conservative approach where the signal had to be twice as high as this noise cut-off (**Fig. S6**, blue dots) in *both replicates* from the same sample. To be "seronegative", the peptide signal needed to be less than this threshold in *both replicates*. Otherwise, the peptide is "not seroreactive". Notably, the seroreactivity thresholds across all plates were in the same range, which reassured us that this approach could be taken for all samples.

**Determination of peptides that are enriched post-exposure**

NHPs: We evaluated the VectorScan repertoires of each NHP separately. To be considered a post-Dengue exposure hit within an NHP, a peptide needed to pass several criteria. First, the pre-exposure sample must be seronegative. Next, to minimize noise, the peptide must either be seroreactive or seronegative at either timepoint. This means that for both timepoints, both replicates must either be above the seroreactivity threshold or below it; there can be no mismatch. Then, to identify hits, we required at least one timepoint to be seroreactive. Finally, to ensure that we analyzed peptides with a robust increase in signal, the difference in signal for the seroconverted timepoint(s) had to be at least 1 (which is the approximate difference between the seroreactivity threshold and the noise threshold for each sample, visualized by the delta between the red and blue dots in **Fig. S6**).

However, in the process of examining the sequences of the arbovirus-derived signal from these NHPs, we observed that a subset of seroconverted hits contained GS-linkers (which are repeat sequences of glycine-serine residues) comprising much of the peptide sequence (>16 amino acids, "GGSGGGSGGGSGGGSG"). These long GS-linkers are an artifact introduced by the *pepsyn* package due to poor sequence quality from the original Uniprot entries. Therefore, we re-analyzed the signal from each NHP, excluding peptides with these long GS-linkers across the entire library, to ensure that we were correcting a systematic error that was introducing non-specific signal, rather than biasing our analyses against any one species. This cleanup decreased the number of peptides being analyzed from 253,789 to 249,253, and had the largest impact on the arboviral section of the library (**Table S1**). None of the random peptide sequences were affected by this filtering, so the seroreactivity thresholds for all samples remained intact.

In addition, to ensure that downstream overrepresentation analyses were being performed using the true number of peptides successfully expressed in the library, we also excluded peptides that had no signal in any replicates in our library characterization (based on the data from sequencing the input library; the non-zero counts were used to generate **Fig. 1C**) from downstream analysis. This further reduced the subset of peptides being analyzed to 249,243 (98.2% of the original library). The enrichment of viral, flaviviral, and DENV signals in each NHP were consistent even after implementing these filtering steps (**Fig. S5**).

NIH: We followed a similar approach to identify peptides that were enriched post-mosquito exposure within each person. Based on the NHP results, we only analyzed signal from peptides if their sequence was not dominated by a GS-linker. Among these downselected peptides, we required that a peptide must be either seroreactive or seronegative at either timepoint (i.e., both replicates needed to be either above or below the threshold for seroreactivity). To ensure that we were only analyzing peptides with a robust increase in signal at Day 44 (the post-exposure timepoint), we set a requirement whereby the difference in signal between Day 0 (the pre-exposure timepoint) and Day 44 had to be at least 1 (which is the approximate difference between the seroreactivity threshold and the noise threshold for each sample, visualized by the delta between the red and blue dots in **Fig. S6**). For a peptide to have "seroconverted," the peptide must be seronegative at Day 0 seroreactive at Day 44. By setting this criterion, we are inherently only looking for antibody responses that arose post-exposure (i.e., *de novo* responses rather than boosting).

**Determination of seroreactive peptides in non-longitudinal (CDC) human samples**

Again, a similar approach to the NIH cohort was taken. The seroreactivity threshold was calculated based on the distribution of random peptides from each sample. A peptide was deemed to be seroreactive if both replicates were above this threshold.

**Comparison of seroprevalence rates among control peptides**

We evaluated the validity of our approach by evaluating the seroprevalence rates of the positive control peptides that were included in VectorScan. These include peptides from common viruses such as rhinovirus B, respiratory syncytial virus, and enterovirus B. We found rates of seroprevalence in the NIH and CDC cohorts that were in line with previous studies (**Table S4**), which gave us confidence in the thresholds that were being set. The same thresholding procedure was applied to the NHP samples, though the seroprevalence of the control peptides was not assessed for those samples since human seroprevalence rates would not be predicted to translate to captive NHPs.

**Network graphing of immunogenic motifs**

NHP Arbovirus hits (Fig. 2F and Fig. S4): For the alphavirus-derived hits with $p<0.01$, a blast database was created (example command: makeblastdb -in Plate22_top_features_Alphavirus.faa -out db -dbtype prot"). *blastp* was then run with the following options: "-evalue 1 -max_hsps 1 -max_target_seqs 100000 -word_size 7 -outfmt 6". Query-target matches with a bit-score ≥ 40 were organized into clusters in *igraph* based on Louvain community detection. Nodes were colored based on the species origin of the peptide, and all clusters (including those with single members) were graphed. The same methodology was used for the flavivirus-derived hits.

Figure 4C: Based on the 3,971 peptides with increased fold change values in at least 2 people, a blast database was created. *blastp* was then run with the following options: "-evalue 1 -max_hsps 1 -max_target_seqs 100000 -word_size 7". Query-target matches with a bit-score ≥ 40 were organized into clusters in *igraph* based on Louvain community detection. Clusters with ≥ 2 members were considered to represent an epitope-containing motif, and clusters with ≥5 members were visualized with a network graph. Nodes were colored based on cluster ID.

Figure 5D: For mosquito-derived peptides with $p<0.01$, a blast database was created and a *blastp* search was performed with the options above. Query-target matches with a bit-score ≥ 40 were organized into clusters in *igraph* based on Louvain community detection. Nodes were colored based on whether the mean peptide signal was higher in the "Known arbovirus exposure" group or the "Unknown exposure" group.

Figure 5F: All peptides that were in clusters of 2 or more members in Fig. 4C and 5D were concatenated into a single fasta file. A blast database was created and *blastp* search was performed with the options above. Query-target matches with a bit-score ≥ 40 were organized into clusters in *igraph* based on Louvain community detection. Nodes were colored based on whether the peptide was in Fig. 4C (NIH) or Fig. 5D (CDC).

**Protein-level domain and functional annotation of mosquito-derived peptides**
Using the sequences from the 11,839 vector-derived proteins that were tiled through to create the vector portion of VectorScan, we performed DIAMOND mapping to identify OrthoMCL entries for each protein. This yielded a set of proteins with 11,614 unique gene ID's, which we downloaded and used as a "Gene ID input set" to query VectorBase

(which is hosted by VeuPathDB). This yielded protein-level annotations for 11,727 entries; this increase in number is because some input genes yielded multiple outputs. We merged these tables to match up each vector-derived protein in VectorScan with any available OrthoMCL group information and protein-level annotations based on the Gene ID. As a result, 11,796 out of the original 11,839 vector-derived proteins had a VectorBase mapping result. After these steps, 2,626 proteins had no associated annotations from VectorBase. While analyzing these results, we observed that some proteins belonged to the same Ortholog group but were missing annotations. We performed some manual annotation of the InterPro annotations for these groups, which is documented in an R Notebook in the GitHub repository accompanying this work. This decreased the number of proteins with no InterPro annotations from 2,939 to 2,812 and the total number of proteins with no annotations to 2,529, or 21% of the original set of vector-derived proteins. We then mapped back these protein-level annotations to each peptide in the library, and 131,494/156,876 (83.9%) of the mosquito-derived peptides had at least one annotation, and 127,612/156,876 had at least 1 InterPro annotation. We then "expanded" this table, so that each row in a table contained all the peptide information, and a single InterPro annotation, effectively creating our database of 503,680 entries.

To identify which InterPro annotations are the most frequent among the peptides that have increased signal in at least two people, we narrowed down our database based on peptide ID (going from 503,680 entries to 11,162). We then created a table of how often each InterPro annotation occurred among these peptides. The "N/A" annotation was the most frequent, but was not included in Fig. 4A due to its uninformative nature. To determine whether each annotation was overrepresented within this set of peptides, we created a contingency table for each annotation.

| | Peptide has annotation X | Peptide does not have annotation X |
|---|---|---|
| Is a hit | X | Y |
| Not a hit | Z | 503,860-X-Y-Z |

A two-sided Fisher's exact test was then performed. Both OR>1 and $p<0.05$ were required in order for an annotation to be considered overrepresented. Again, the "N/A" annotation had the highest overrepresentation, which is typical of GO analysis with non-model organisms, but was excluded from analysis (i.e., Fig. 4B) because of the lack of information that such an annotation provides.

### Supplementary Figures

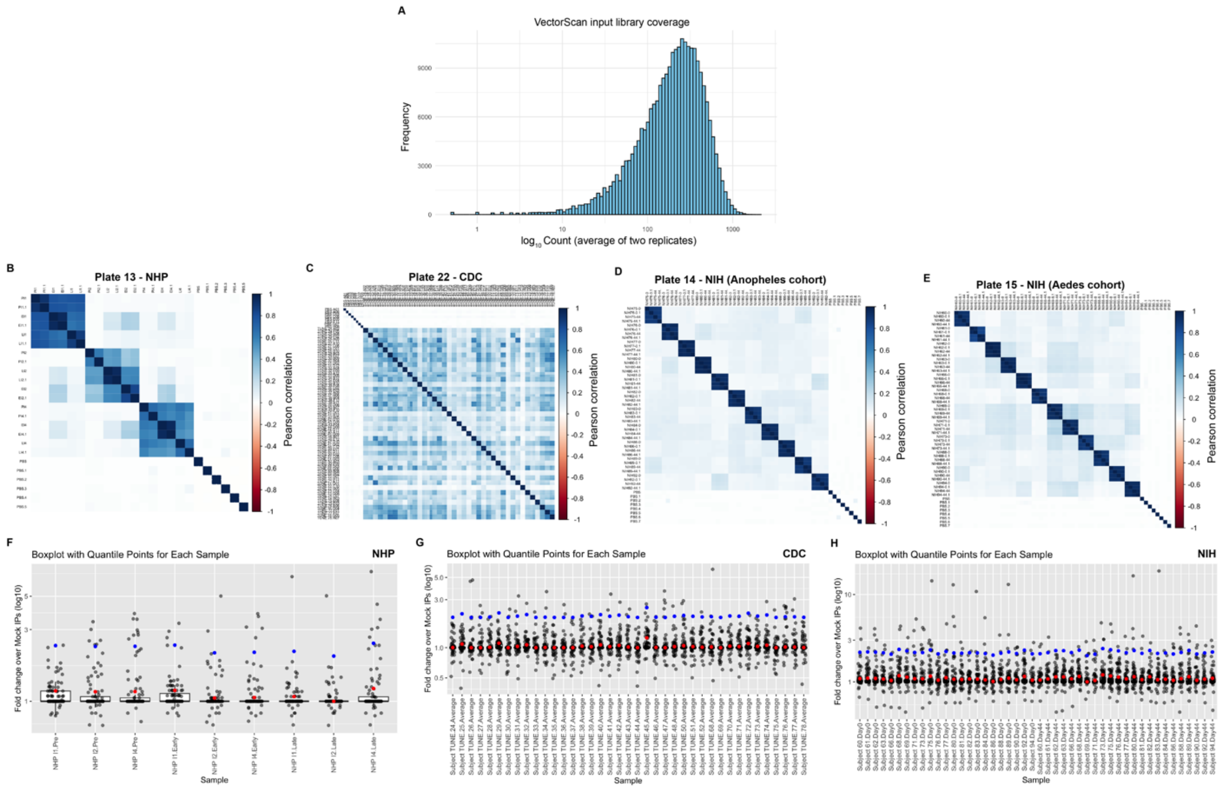

**Figure S1: Library and sample characterization via correlation and fold change (FC) analyses. (A)** The expanded library was sequenced to a read depth of approximately 250 times the size of the library. Counts across two replicates were averaged. 612 peptides had 1 or fewer reads (peptides with 0 reads are not represented due to the log10 x-axis), indicating 99.76% library coverage. 77.1% of peptides had an average read count between 100 and 1000. **(B-E)** Pearson correlation coefficients were calculated for each pair of samples within each plate that was sequenced. The fold change values for each replicate across all 253,789 peptides in the library were included in the calculation. High correlation between replicates and low correlation among PBS controls was used as a quality control measure. **(F-H)** Fold change distributions among the 100 random peptides for each cohort of samples.

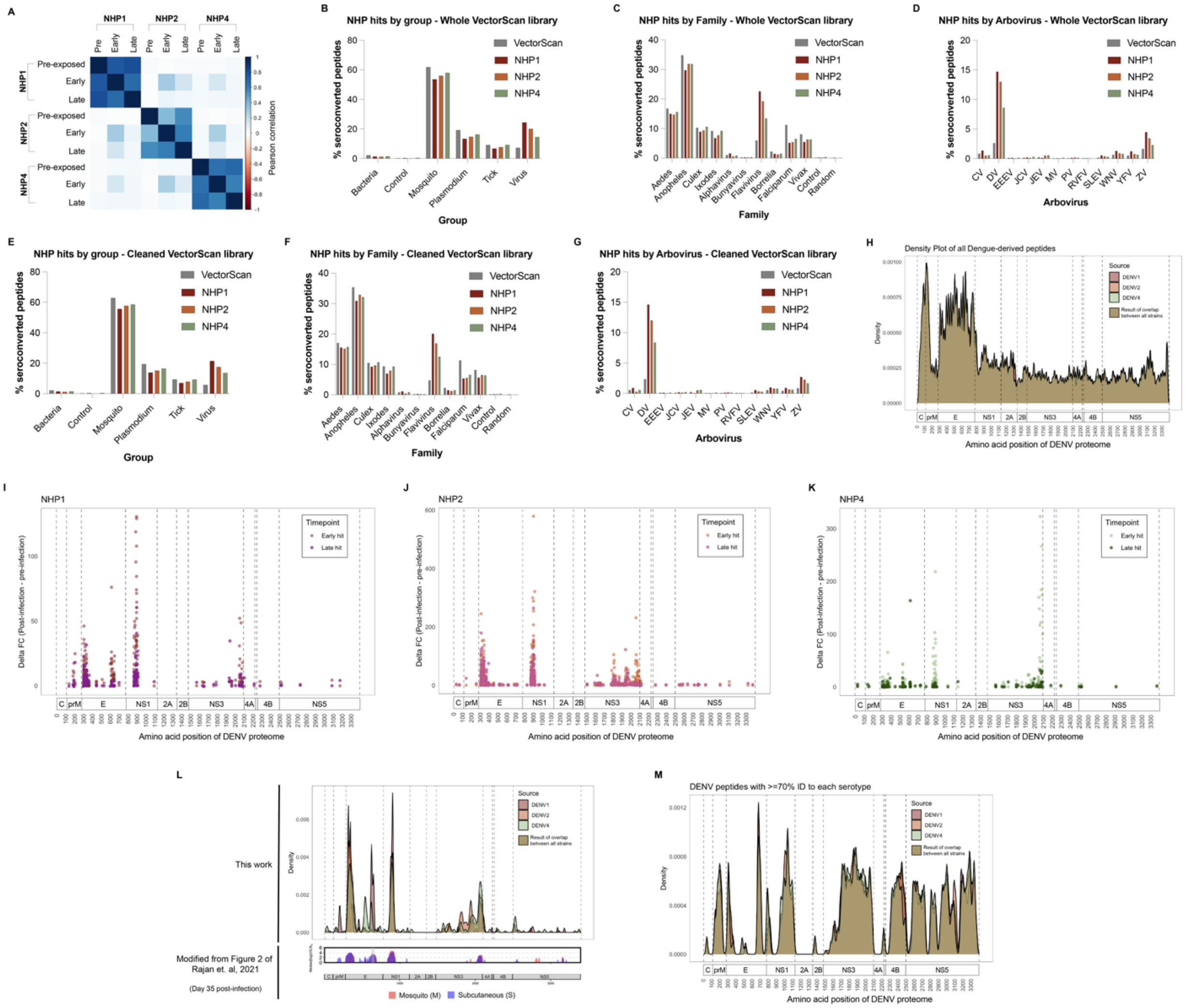

**Figure S2: Analysis of VectorScan profiles from DENV-infected NHPs, with a focus on DENV-derived peptides. (A)** The Pearson coefficient between each pair of samples was calculated based on transformed read counts (normalized against serum-free controls and averaged across replicates) from each peptide. **(B-G)** Peptide categorization is by general group, family, or arbovirus species. **(B-D)** Percentages are based on representation across all 253,789 peptides in the VectorScan library. **(E-G)** Percentages are based on the "cleaned" version of the library (249,243 peptides), after excluding peptides with long GS-linkers and those that were not expressed based on library characterization with high sequencing depth. **(H)** All DENV-derived peptides from the cleaned library were aligned to the DENV1, DENV2, and DENV4 strains that were used to infect the NHPs. The amino acid position of the peptide is based on the midpoint of the *blastp* alignment. **(I-K)** The difference between the average fold change values at timepoint X and the pre-exposure timepoint was calculated for each seroconverted peptide. **(L)** A previous study using a phage library focused on flaviviruses profiled samples from NHPs that were infected with DENV1 delivered via subcutaneous (S) injection or mosquitos (M).[13] In this study, significantly enriched peptides were mapped to the DENV1 proteome. Heat maps and the corresponding density plot show the difference in signal between Day 35 post-exposure and baseline. These DENV1-specific signals are plotted beneath the density plots from peptides that seroconverted across all NHPs and at all timepoints in this study. **(M)** Peptides with at least 70% identity across the entire peptide to each DENV strain, indicating regions of the proteome that are most likely to elicit cross-reactive antibodies.

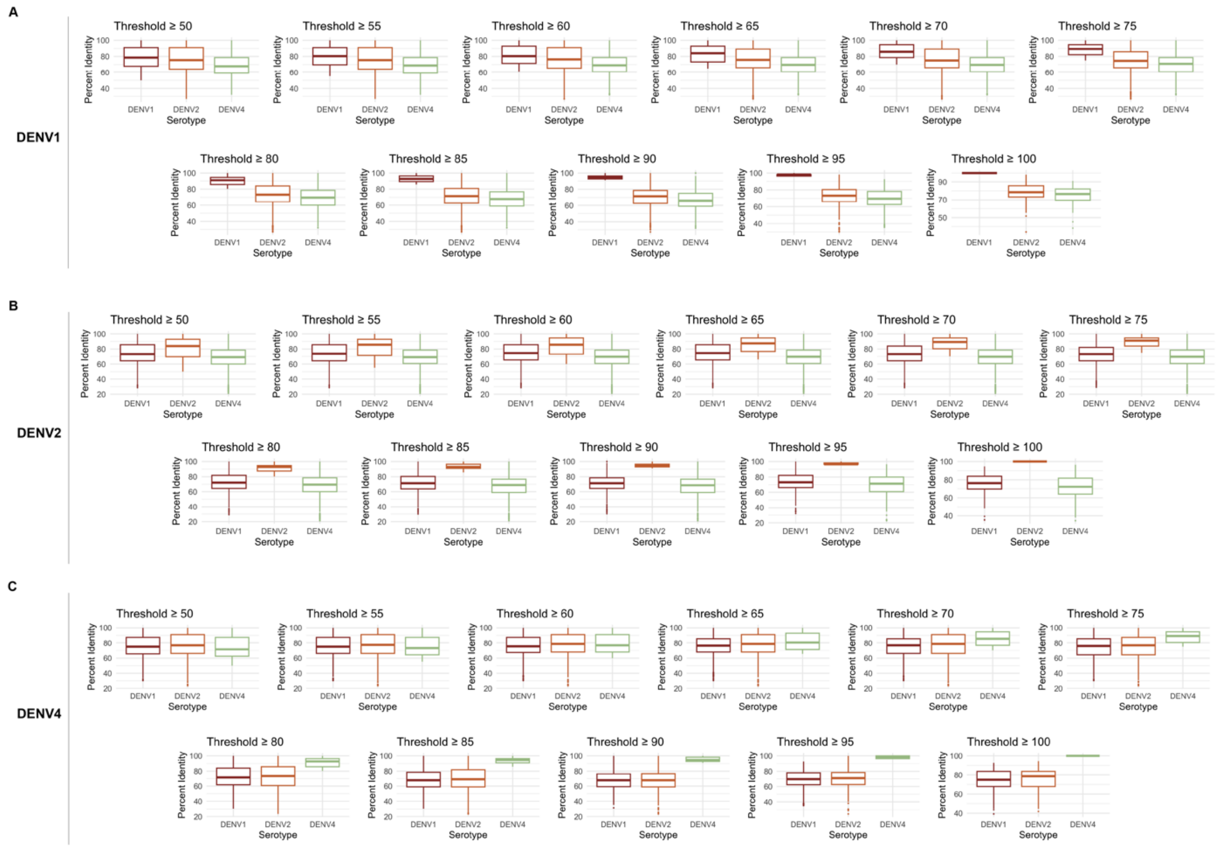

**Figure S3: Benchmarking analysis to define serotype-informative peptides.** Applying increasing thresholds of percent identity to **(A)** DENV1, **(B)** DENV2, and **(C)** DENV4 yields natural separation of peptides based on serotype-specificity.

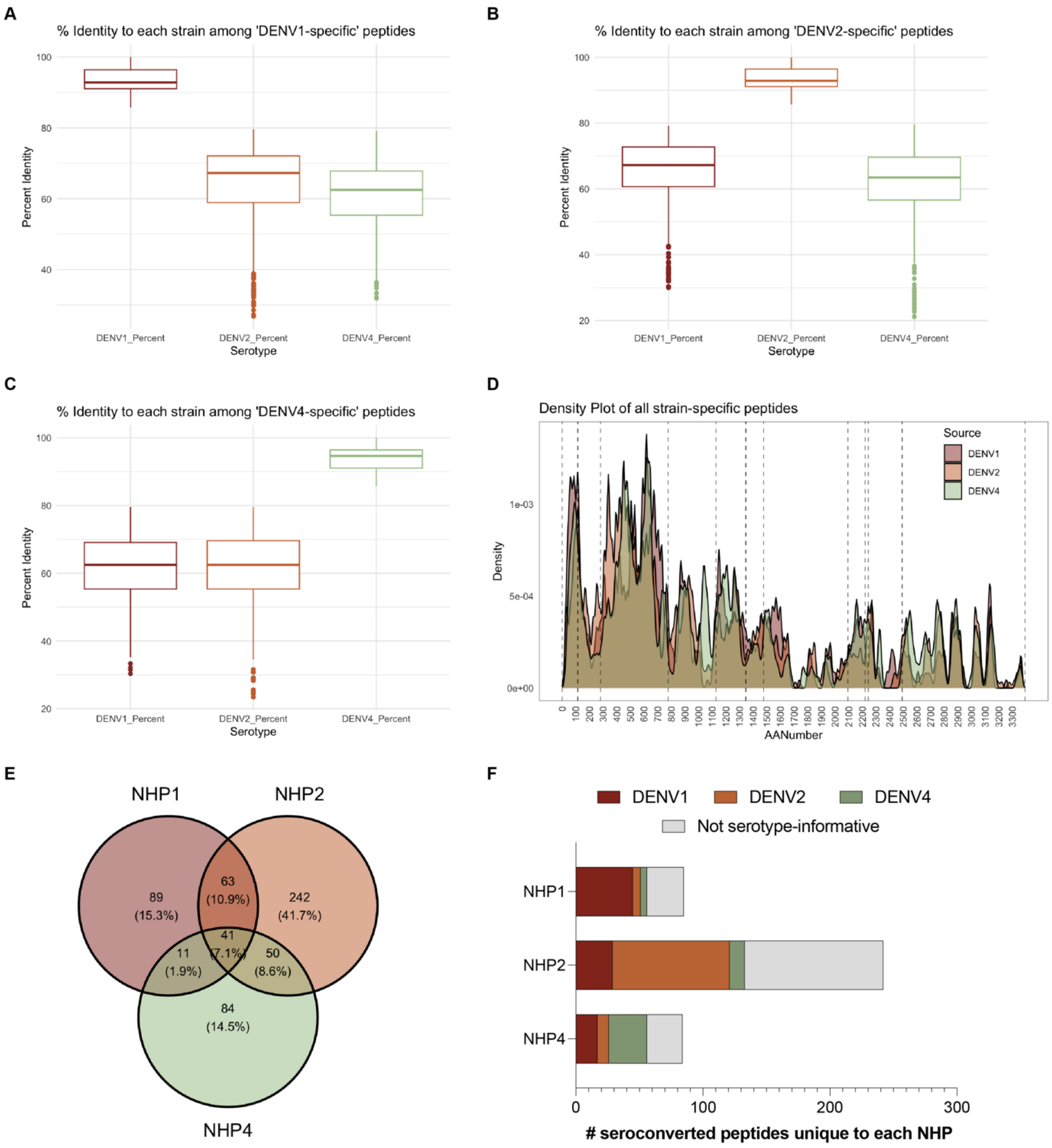

**Figure S4: Identification of peptides and serotype-informative peptides with increased seroreactivity in each animal.** Serotype-informative peptides were determined based on whether (1) there was a full-length peptide match, (2) at least 85% sequence identity to one strain, and (3) less than 80% identity to the other two. **(A-C)** The resulting distributions of sequence identity among these serotype-informative peptides yield a high percentage of identity to the strain of interest, and a median of less than 70% identity to the other two strains. **(D)** These serotype-informative peptides are distributed throughout the DENV proteome. **(E)** The subset of peptides that had post-exposure signal were compared across NHPs. **(F)** Each peptide that was unique to a particular NHP was annotated based on whether it fit the criteria as “serotype-informative”. Peptides that meet the criteria were labeled with the relevant DENV serotype.

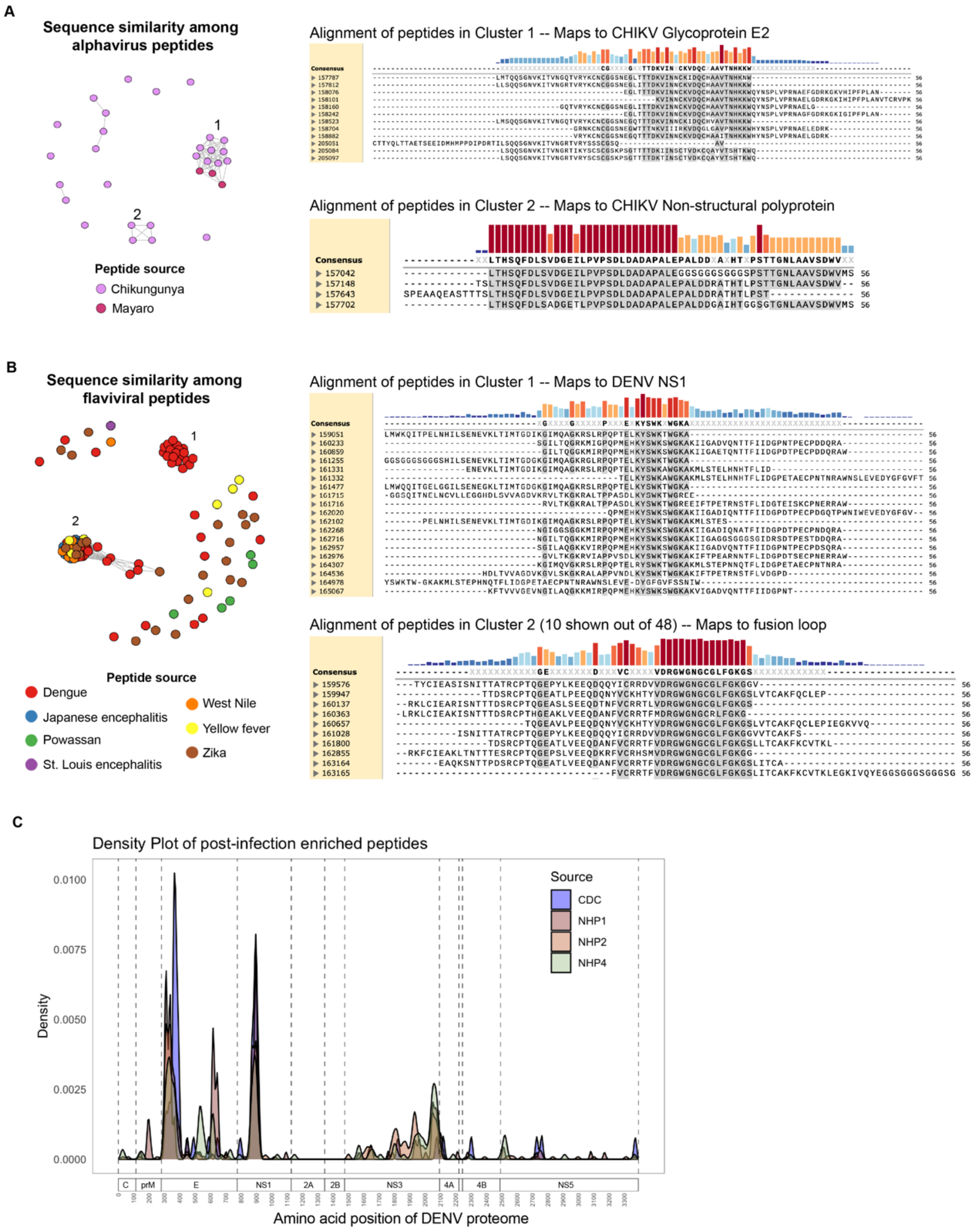

**Figure S5: Sequence-based clustering of peptides that are associated with samples with known arbovirus exposure.** A network graph was created for the **(A)** alphavirus-derived peptides and **(B)** flavivirus-derived peptides. Sequence alignments of peptides from two of the clusters show the shared immunogenic epitope that is driving clustering. Alignments were done via MUSCLE in SnapGene. Positions with at least 70% conservation were used to generate a consensus sequence. The amino acids matching the consensus sequence are highlighted in gray. **(C)** The DENV-derived peptide hits from each NHP and from the set of 28 DENV-derived peptides that were associated with arbovirus exposure in the CDC cohort were mapped to the DENV proteome (the CDC hits were mapped to DENV1).

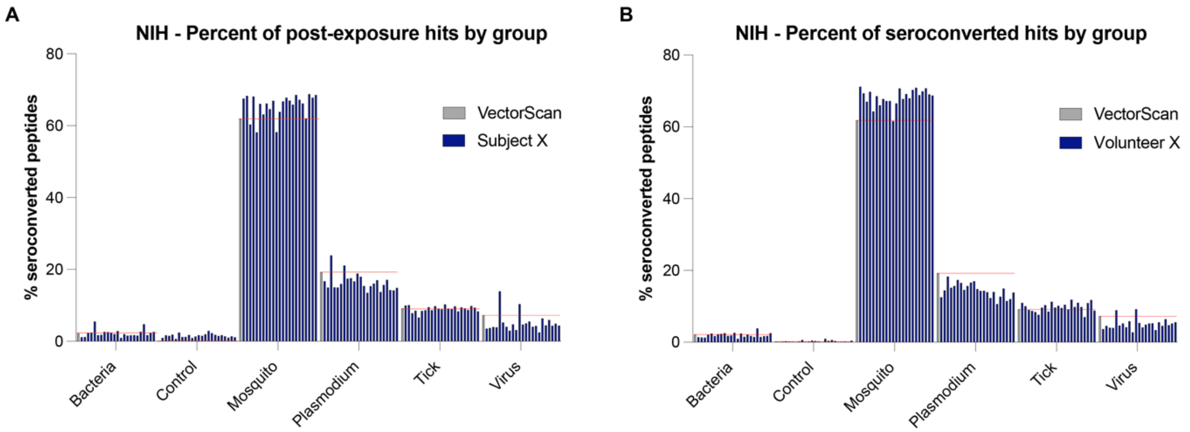

**Figure S6: Percentage of seroconverted peptides from each category in VectorScan among individuals. (A)** Representation among all peptides with signal that is increased at Day 44. **(B)** Representation among all peptides that seroconverted between Day 0 and Day 44. Red lines mark the percentage of representation of each group in the cleaned VectorScan library (gray bars), to assist with visual analysis of population- and individual-level (blue bars) overrepresentation.

### Supplementary Tables

**Table S1: Protein sources for organisms included in VectorScan and counts of their final peptide representation.** Contains the Uniprot taxonomy ID that was used to compile the list of protein sequences ("No. starting sequences") that were subsequently collapsed based on identity or downselected based on desired features and processed for peptide tiling. Peptide counts from the different genera/species represented in VectorScan, before ("Original Library") and after filtering out peptides with long GS-linkers ("Cleaned").

**Table S2: VectorScan library sequences.** Sequences for all 253,789 peptides, and associated source information.

**Table S3: Clusters of peptides resulting from epitope analysis of the NIH and CDC cohorts**.

**Table S4: Control seroprevalence rates**. Comparisons of seroprevalence rates from the human samples screened by VectorScan for control peptides.
